# Impaired dendritic spike generation in the Fragile X prefrontal cortex is due to loss of dendritic sodium channels

**DOI:** 10.1101/2022.05.11.491501

**Authors:** Federico Brandalise, Brian E. Kalmbach, Erik P. Cook, Darrin H. Brager

## Abstract

Patients with Fragile X syndrome, the leading monogenetic cause of autism, suffer from impairments related to the prefrontal cortex including working memory and attention. Synaptic inputs to the distal dendrites of layer 5 pyramidal neurons in the prefrontal cortex have a weak influence on the somatic membrane potential. To overcome this filtering, distal inputs are transformed into local dendritic Na^+^ spikes, which propagate to the soma and trigger action potential output. Layer 5 extratelencephalic (ET) PFC neurons project to the brainstem and various thalamic nuclei and are therefore well positioned to integrate task-relevant sensory signals and guide motor actions. We used current clamp and outside-out patch clamp recording to investigate dendritic spike generation in ET neurons from male wild type and *Fmr1* knockout (FX) mice. The threshold for dendritic spikes was more depolarized in FX neurons compared to wild type. Analysis of voltage responses to simulated in vivo “noisy” current injections showed that a larger dendritic input stimulus was required to elicit dendritic spikes in FX ET dendrites compared wild type. Patch clamp recordings revealed that the dendritic Na^+^ conductance was significantly smaller in FX ET dendrites. Taken together, our results suggest that input-output transformation is impaired in ET neurons of the PFC in FX mice. Considering our prior findings that somatic D-type K^+^ and dendritic HCN-channel function is reduced in ET neurons, we suggest that the integration of information by PFC circuits is fundamentally altered in Fragile X syndrome.

**KEY POINTS:**

- Dendritic spike threshold is depolarized in Layer 5 PFC neurons in FX mice
- Simultaneous somatic and dendritic recording with white noise current injections revealed that larger dendritic stimuli were required to elicit dendritic spikes in FX ET neurons
- Outside-out patch clamp recording revealed that dendritic sodium conductance density was lower in FX ET neurons

## INTRODUCTION

The prefrontal cortex (PFC) is involved in working memory tasks and contributes to goal-directed behavior by selecting and executing task-appropriate responses while suppressing maladaptive or task-inappropriate ones (Goldman-Rakic, 1990; Rainer *et al*., 1998; Miller & Cohen, 2001; Sakai *et al*., 2002; Lara & Wallis, 2015). The PFC sits at the top of the hierarchy associated with cognitive function, by exerting top-down control over numerous cortical and subcortical regions (Kesner & Churchwell, 2011). Output from the deep layers, is carried by the axons of both extratelencephalic (ET) and intratelencephalic (IT) pyramidal neurons (Molnár & Cheung, 2006; Shepherd, 2013; Baker *et al*., 2018). These neuron types differ in morphology, response to neuromodulation, distribution of ion channels and long-range targets (Dembrow & Johnston, 2014). L5 ET neurons project to the brainstem and various thalamic nuclei and are therefore well positioned to integrate task-relevant sensory signals and guide motor actions. Higher order thalamic nuclei receive feedforward inputs from L5 ET neurons of the PFC (Li *et al*., 2003; Sherman, 2016). Thalamic nuclei in turn send connections to the superficial layers (L1) of the PFC (Groenewegen, 1988; Deniau *et al*., 1994). Evidence suggests that there are strong reciprocal connections between the PFC and thalamic nuclei and that these thalamocortical loops are critical for cognitive processing (Schmitt *et al*., 2017; Collins *et al*., 2018).

In cortical neurons, the distal dendrites, where the majority of synaptic inputs impinge, and the axosomatic region, where action potentials are generated, are electrically isolated from each other due to both neuronal morphology and the presence of voltage-gated ion channels including h-channels (Berger *et al*., 2003; Harnett *et al*., 2013; Dembrow *et al*., 2015). In contrast to L5 neurons in sensory cortex, which rely on Ca^2+^ plateaus (Larkum *et al*., 1999; Larkum *et al*., 2003), synaptic inputs in the distal dendrites of L5 ET PFC neurons are transformed into local dendritic Na^+^ spikes, which can propagate to the soma and result in action potential output (Remy *et al*., 2009; Kalmbach *et al*., 2017). These dendritic nonlinearities are believed to play important roles for neuronal computations in cortical circuits (Poirazi *et al*., 2003*a*; 2003*b*; London & Häusser, 2005; Ujfalussy *et al*., 2015; Kaifosh & Losonczy, 2016).

Patients with Fragile X syndrome (FXS), the most common form of inherited cognitive impairment and leading monogenic cause of autism, suffer deficits related to PFC function including: attention and working memory deficits, impulsivity and behavioral inflexibility (Munir *et al*., 2000*a*; 2000*b*; Wilding *et al*., 2002; Bray *et al*., 2011). In rodents, sensorimotor coordination has been investigated using a number of behavioral tasks (Siegel *et al*., 2012; Guo *et al*., 2014; Murakami *et al*., 2014; Goard *et al*., 2016). In many of these behavioral tasks, when the cue is separated in time from the stimulus, the task becomes PFC-dependent and assumes the structure of a working memory task. The prevailing hypothesis is that a persistent signal to downstream brain regions is necessary to bridge the temporal gap between cue and stimulus (Weiss & Disterhoft, 2011; Siegel *et al*., 2012; Guo *et al*., 2017; Economo *et al*., 2018). L5 ET neurons, which project directly to thalamus and other downstream brain regions, are a prime candidate for providing this persistent signal and modulate many of the PFC-dependent behavioral phenotypes associated with FXS (Nakayama *et al*., 2018).

We recently showed that dendritic spike generation is impaired in CA1 pyramidal neurons of FX mice (Ordemann *et al*., 2021). In the present study, we investigated dendritic spike generation in ET neurons of FX mice. We found that the threshold for dendritic spikes was more depolarized, and the maximum rate of rise was slower for dendritic spikes in FX ET dendrites compared wild type. Analysis of outside-out patch clamp recordings revealed the density of dendritic voltage-gated Na^+^ channels was significantly lower in FX neurons compared wild type. The net consequences of these changes impact the ability of ET neurons of the PFC to reliably transform synaptic inputs into action potential output with potential negative impacts on PFC circuit function in Fragile X syndrome.

## Materials and Methods

### Ethical Approval

All procedures involving mice were performed with the approval of the University of Texas Institutional Animal Care and Use Committee. All wild type and Fmr1 knockout mice (C57Bl/6 male, 2-4 months old) were housed in the satellite animal facility under the management of the University of Texas Animal Resource Center and in reversed light/dark cycles.

### Acute slice preparation

Mice were anesthetized with a ketamine/xylazine (100 mg/kg, 10 mg/kg) cocktail and were perfused through the heart with ice-cold saline consisting of (in mM): 2.5 KCl, 1.25 NaH_2_PO_4_, 25 NaHCO_3_, 0.5 CaCl_2_, 7 MgCl_2_, 7 dextrose, 205 sucrose, 1.3 ascorbate and 3 sodium pyruvate (bubbled with 95% O_2_/5% CO_2_ to maintain pH at ~7.4). A vibrating tissue slicer (Vibratome 3000, Vibratome Inc.) was used to make 300 μm thick coronal sections containing the prefrontal cortex. Slices were held for 30 minutes at 35°C in a chamber filled with artificial cerebral spinal fluid (aCSF) consisting of (in mM): 125 NaCl, 2.5 KCl, 1.25 NaH_2_PO_4_, 25 NaHCO_3_, 2 CaCl_2_, 2 MgCl_2_, 10 dextrose and 3 sodium pyruvate (bubbled with 95% O_2_/5% CO_2_) and then at room temperature until the time of recording. Note that wild type and FX mice were interleaved daily, when possible, although the experimenter was not blind to the genotype.

### Electrophysiology

Slices were placed in a submerged, heated (32–34°C) recording chamber that was continually perfused (1−2 ml/minute) with bubbled aCSF containing (in mM): 125 NaCl, 3.0 KCl, 1.25 NaH_2_PO_4_, 25 NaHCO_3_, 2 CaCl_2_, 1 MgCl_2_, 10 dextrose, 3 sodium pyruvate. Slices were viewed either with a Zeiss Axioskop microscope and differential interference optics. Patch pipettes (4−8 MΩ) were pulled from borosilicate glass and wrapped with Parafilm to reduce capacitance. All recordings were made from L5 ET pyramidal neurons in the anterior cingulate or prelimbic regions of mPFC.

### Whole cell recordings

The pipette solution contained (in mM): 120 K-gluconate, 16 KCl, 10 HEPES, 8 NaCl, 7 K_2_ phosphocreatine, 0.3 Na−GTP, 4 Mg−ATP (pH 7.3 with KOH). Neurobiotin (Vector Laboratories; 0.1-0.2%) was also included for histological processing and post-hoc cell location determination. In some experiments, FMRP_1-298_ (100 nM; Novus Biologicals H00002332-01) or heat-inactivated (90°C for 10 min) FMRP_1-298_ was included in the internal recording solution. Alexa 594 (16 μM; Thermo Fisher #A10428) was also included in the internal recording solution to determine the dendritic recording location relative to the soma. Data were acquired using a Dagan BVC−700 amplifier (Dagan Inc.) and custom data acquisition software written using Igor Pro (Wavemetrics). Data were acquired at 10−50 kHz, filtered at 2−10 kHz and digitized by an ITC-18 (InstruTech) interface. Pipette capacitance was compensated for, and the bridge was balanced during each recording. Series resistance was monitored and compensated throughout each experiment and was 10−25 MΩ for somatic recordings and 15-40 MΩ for dendritic recordings. Recordings were discarded if series resistance increased by more than 30% during the recording. Voltages are not corrected for the liquid-junction potential (estimated as ~12 mV).

### White noise injections and Leaky-integrate-and-fire model

Dendritic recordings were performed with a large, zero-mean, Gaussian-distributed white-noise stimulus current injected through the recording pipette for 1–2 min. The variance of the noise was adjusted to two levels: large, to produce somatic action potentials and small, subthreshold for somatic actin potentials. We implemented a leaky-integrate-and-fire (leaky IAF) model to simulate the effects of a change in dendritic spike threshold on dendritic integration. The leaky integrator was convolved with Gaussian distributed white-noise current injections created using the same standard deviations as that used in the actual recordings. Thus, the WT and FX models received the same statistical white-noise inputs as the WT and FX experiments, respectively. The leaky integrator was modeled as a sum of two exponentials with decay constants = 0.5 and 1.8 ms, weighted by 1.1 and 1.0, respectively (shown in Fig. 4E), and was the same for all models. Spike thresholds were set to produce enough spikes required for the STAs (> 1000 spikes). Thresholds were initially set to 7 for both the WT and FX models, and then the FX model ‘s threshold was increased to 11.5. When the convolution of the leaky integrator and the white-noise stimulus (*x* in Fig. 4E) crossed the threshold (*T*), the model produced a spike, and the leaky integrator was reset to zero. The model ‘s spike times were randomly jittered using a Gaussian distribution with a standard deviation of 100 us to mimic recording noise.

### Outside-out recordings

Outside-out recordings were made using an Axopatch 200B amplifier (Molecular Devices), sampled at 10 kHz, analog filtered at 2 kHz and digitized by an ITC-18 interface connected to a computer running Axograph X. The pipette solution contained (in mM): 90 K-gluconate, 50 CsCl, 10 HEPES, 5 EGTA, 4 NaCl, 7 K_2_ phosphocreatine, 0.3 Na−GTP, 4 Mg−ATP (pH 7.3 with KOH). TEA-Cl (20) and 3,4 diaminopyridine (0.1 mM) were added to the extracellular saline. I_Na_ was measured using depolarizing voltage commands from −80 to 40 mV from a holding potential of −90 mV. Patch area was estimated by fitting the decay of the capacitive transient in response to a small voltage step (assuming 1 µF cm^-2^; Routh *et al*., 2017). Activation data were fit to a single Boltzmann function using a least-squares program. Linear leakage and capacitive currents were digitally subtracted by scaling traces at smaller command voltages in which no voltage-dependent current was activated.

### Statistical Analysis

Repeated measures analysis of variance (RM−ANOVA), between-subjects factors ANOVA, mixed factors ANOVA and post−hoc t−tests were used to test for statistical differences between experimental conditions. Sidak ‘s correction was used to correct for multiple comparisons. Pearson ‘s product moment correlation was used to test for statistically significant correlations between variables. Data are presented as mean and error bars represent standard error of measurement (SEM). Statistical analyses were performed using Prism (Graphpad) and considered significant if p<0.05. Power analyses were performed using G*power and reported as Type II error probability (β).

## RESULTS

### Dendritic spike generation is impaired in FX ET neurons

In many pyramidal neurons, distal dendritic voltage signals are strongly attenuated by the time they reach the soma. L5 ET neurons generate dendritic Na^+^ spikes to overcome the electrotonic attenuation of distal synaptic inputs (Dembrow *et al*., 2015; Kalmbach *et al*., 2017). We first compared dendritic spikes between wild type and FX ET neurons using dendritic recording and square pulse depolarizing injections (Fig. 1A). The voltage threshold for dendritic spikes was calculated as 20% of the second peak of the second derivative as previously described (Gasparini *et al*., 2004; Ordemann *et al*., 2021). The threshold for dendritic spikes did not vary with distance from the soma in either wild type or FX ET neurons (Fig. 1B). Dendritic spike threshold was, however, significantly more depolarized in FX compared wild type dendrites (wild type: −30.2 ± 0.69 mV, n=32; FX: −24.8 ± 1.35 mV, n=21; unpaired t-test: t=3.869, p = 0.003;). We grouped dendritic spikes into proximal (<250 µm) and distal (>250 µm, near the nexus) based on the distance of the recording electrode from the soma. Threshold was more depolarized for both proximal and distal dendritic spikes (Fig. 1C; wild type: proximal n = 10, distal n = 22; FX: proximal n = 11, distal n = 10; two-way ANOVA, main effect of genotype: F_(1, 49)_=13.26, p = 0.0007). The maximum rate of rise and peak membrane potential for dendritic spikes decreased with distance from the soma for both wild type and FX ET dendrites (Fig. 1D, F). The maximum rate of rise was significantly slower in FX dendrites compared to wild type dendrites (Fig. 1E; wild type: proximal n = 10, distal n = 22; FX: proximal n = 7, distal n = 10; two-way ANOVA, main effect of genotype: F_(1, 44)_=14.31, p = 0.0005). By contrast, there was no significant difference in peak membrane potential between FX and wild type dendritic spikes (Fig. 1G; wild type: proximal n = 12, distal n = 26; FX: proximal n = 13, distal n = 10; two-way ANOVA, main effect of genotype: F_(1, 57)_=0.3475, p = 0.5579).

**Figure 1.**
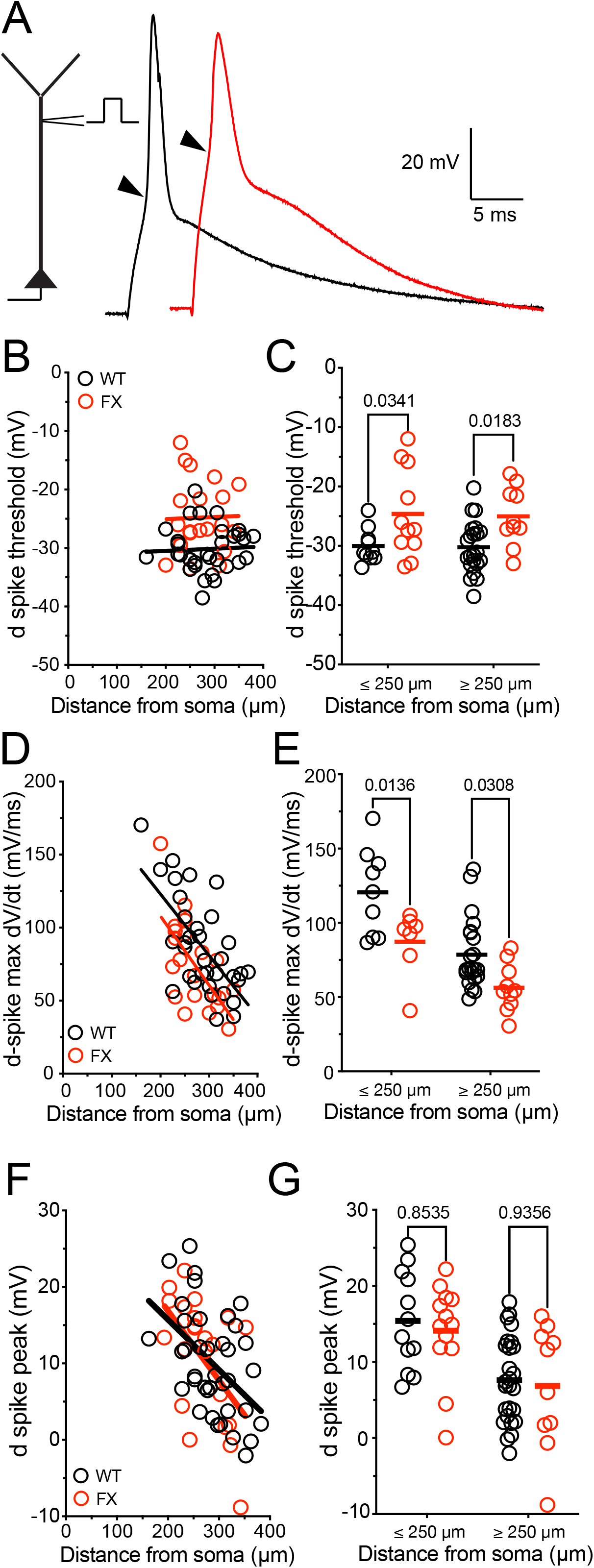
Dendritic spikes are altered in FX ET neurons. A, Representative dendritic spikes recorded from wild type (black) and FX (red) ET dendrites. Threshold is indicated by the black arrow heads. B, Dendritic spike threshold as a function of dendritic location for wild type and FX dendrites. C, Dendritic spike threshold is significantly more depolarized in FX ET neurons compared to wild type ET neurons. D, Dendritic spike rate of rise (dV/dt) as a function of dendritic location for wild type and FX dendrites. E, Dendritic spike maximum dV/dt is significantly smaller in FX ET neurons compared to wild type ET neurons. F, Dendritic spike peak voltage as a function of dendritic location for wild type and FX dendrites. G, There is no significant difference in dendritic spike peak voltage between wild type and FX ET neurons.

### Back propagating action potentials are smaller in FX ET dendrites

To determine if the impairment of active events was limited to dendritic spikes, we measured the backpropagation of action potentials. We made dendritic current clamp recordings and elicited action potentials in wild type and FX ET neurons using extracellular stimulation in layer 5 in the presence of synaptic blockers to prevent fast glutamatergic and GABAergic transmission (Fig. 2A). We found that the peak membrane potential of backpropagating action potentials (bAP) decreased with increasing distance for both wild type and FX neurons (Fig. 2B). When recording location was binned by distance (proximal: <250 µm; distal: >250 µm), we found that the peak membrane potential of distal, but not proximal, bAPs was significantly smaller in FX neurons compared to wild type (Fig. 2C; two-way ANOVA, interaction: F_(1, 76)_=4.363, p = 0.0401; distal wild type: 5.3 ± 1.63 mV, n=29; distal FX: −4.4 ± 2.85 mV, n=16). The half width of bAPs increased with distance for both wild type and FX neurons and was significantly longer in FX neurons at distal, but not proximal, recording locations (Fig. 2D-E; two-way ANOVA, interaction: F_(1, 76)_=5.756, p = 0.0189; distal wild type: 1.7 ± 0.121 ms, n=29; distal FX: 2.4 ± 0.219 ms, n=16).

**Figure 2.**
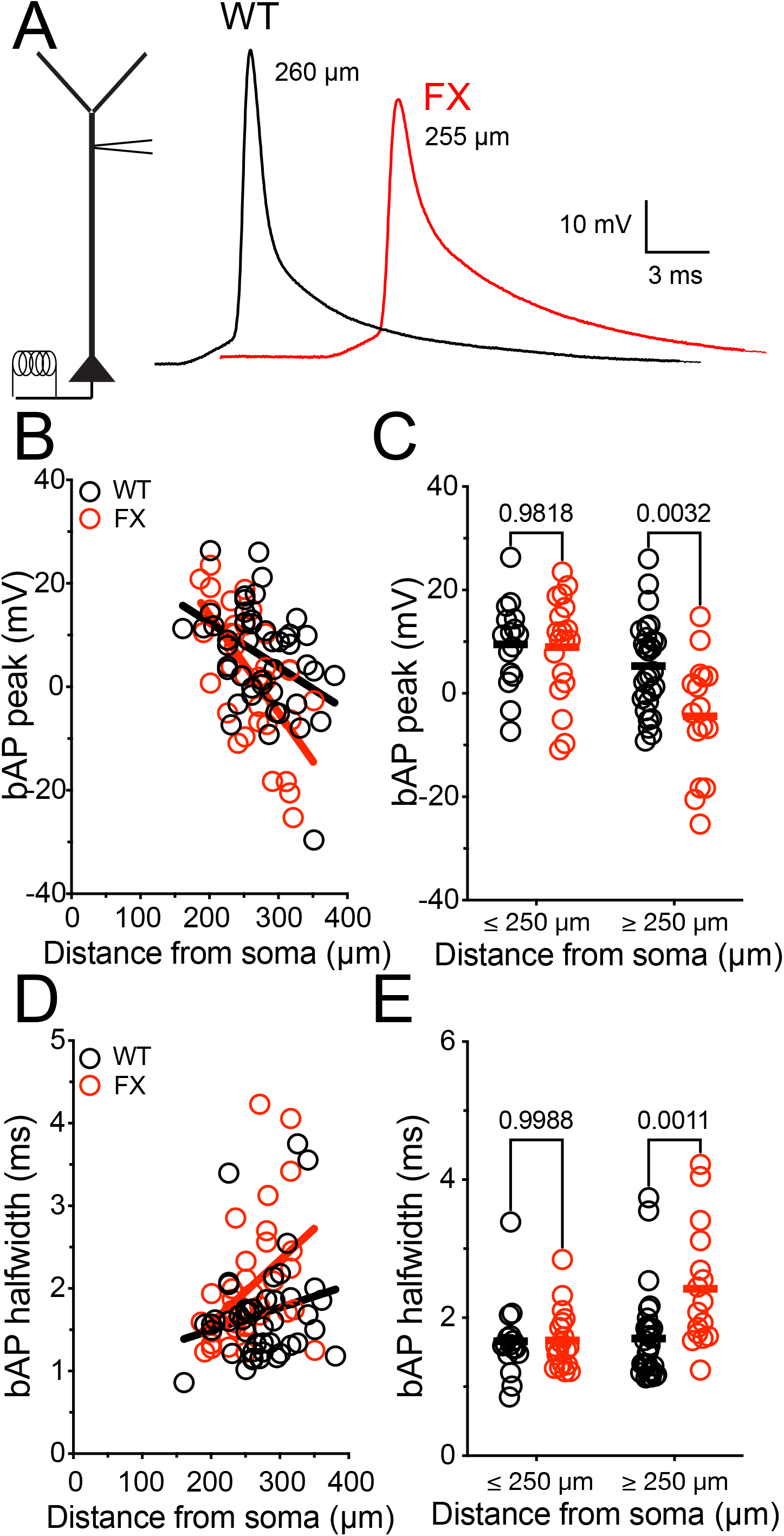
Backpropagating action potentials in the distal dendrites of FX ET neurons are shorter and wider. A, Representative backpropagating action potential recorded from wild type (black) and FX (red) ET dendrites. B, Backpropagating action potential peak as a function of dendritic location for wild type and FX dendrites. C, Backpropagating action potential peak is significantly smaller at more distal locations in FX ET neurons compared to wild type ET neurons. D, Backpropagating action potential halfwidth as a function of dendritic location for wild type and FX dendrites. E, Backpropagating action potential halfwidth is significantly larger at more distal locations in FX ET neurons compared to wild type ET neurons.

### Dendro-somatic coupling is reduced in FX ET neurons

We used simultaneous somatic and dendritic current clamp recordings to investigate the coupling between the soma and dendrites in wild type and FX ET neurons. We used somatic current injection to elicit single action potentials and measure their backpropagation into the dendrites (Fig. 3A). The peak voltage of action potentials in the soma was not significantly different between wild type and FX neurons (Fig. 3B; wild type: 58 ± 3.3 mV, n=6; FX: 57 ± 3.7 mV, n = 7; unpaired t-test: t=0.2951, p =0.7734). In agreement with our results using extracellular stimulation, attenuation of bAPs was significantly greater in FX ET neurons compared to wild type (Fig. 3C). To investigate the ability of dendritic spikes to influence somatic membrane potential, we simulated synaptic input by using double exponential dendritic current injections (Fig. 3D). In agreement with our results using square pulses (Fig. 1), the threshold for dendritic spikes was significantly more depolarized in FX ET neurons compared to wild type (Fig. 3E; wild type: −29 ± 1.6 mV; FX: −12 ± 1.9 mV; unpaired t-test: t=7.023, p =0.001). Furthermore, while dendritic spikes in wild type ET neurons reliably triggered somatic action potentials (median probability = 1.0), this coupling was significantly reduced in FX ET neurons (median probability = 0.8; Fig. 3F; Mann-Whitney test: U=4, p =0.008).

**Figure 3.**
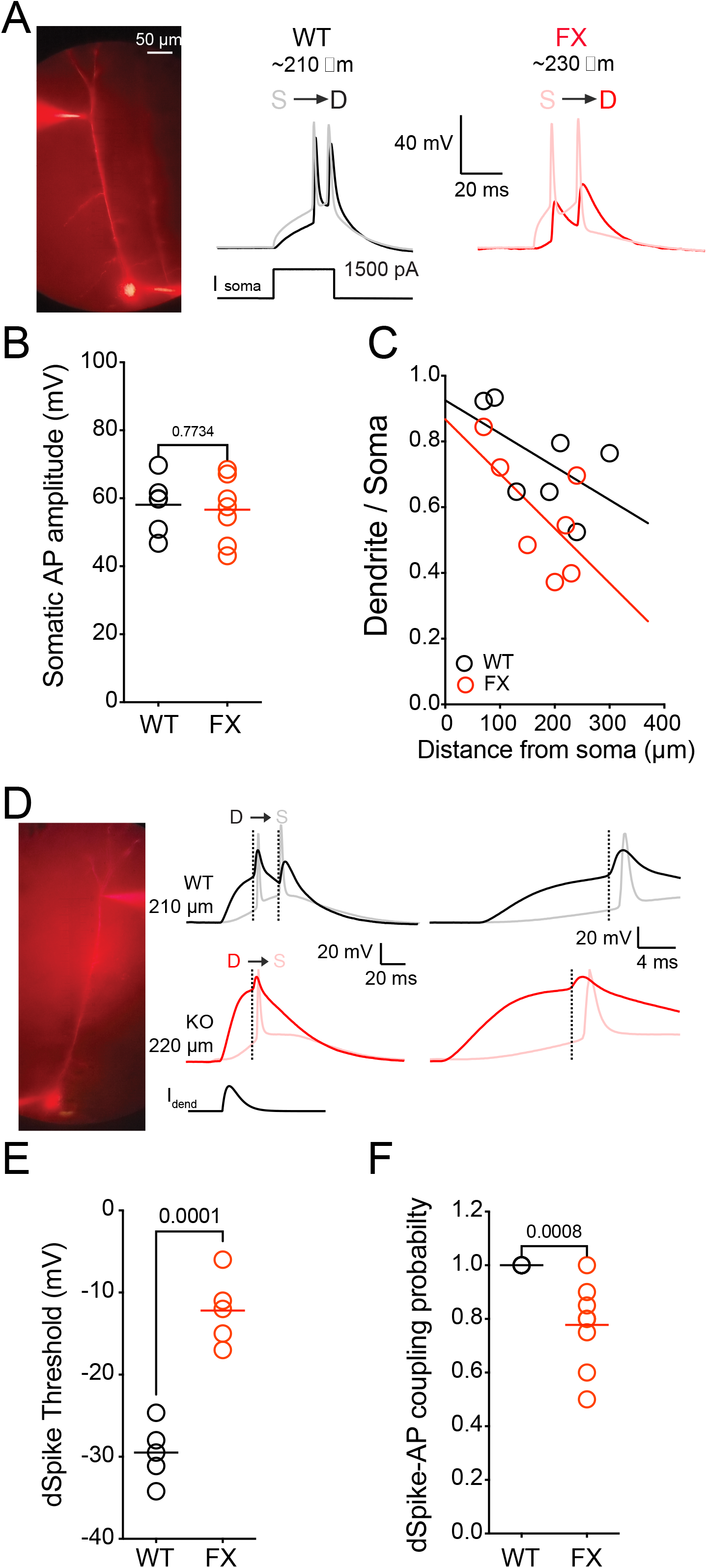
Dendritic-somatic coupling is reduced in FX ET neurons. A, Representative dual somatic-dendritic recording showing dendritic and somatic membrane potential during *somatic* current injection to trigger action potentials in wild type and FX ET neurons. B, Somatic action potential amplitude was not significantly different between wild type and FX ET neurons. C, Attenuation of backpropagating action potentials during dual recording is greater in FX compared to wild type ET neurons. D, Representative dual somatic-dendritic recording showing dendritic and somatic membrane potential during *dendritic* current injection to trigger dendritic spikes in wild type and FX ET neurons. E, Dendritic spike threshold during dual recordings was significantly more depolarized in FX ET neurons compared to wild type. F, The probability of dendritic spikes triggering somatic action potentials was smaller in FX ET neurons.

### Larger stimuli are required to trigger dendritic spikes under in vivo like conditions

To better capture the range of frequencies experienced by the dendrites *in vivo*, we delivered a white-noise stimulus through the dendritic electrode during simultaneous somatic and dendritic recording (Fig. 4A; Kalmbach *et al*., 2017). During the white-noise current injection, it was not possible to reliably identify dendritic spikes present in the dendritic voltage waveform. Thus, we used a previously published systems-based approach that deconvolved out the nonlinear current produced by the dendrites in response to the white noise injection (Kalmbach *et al*., 2017). The deconvolution process revealed inward “current spikes” necessary to account for the dendritic potential. It was previously demonstrated that these dendritic current spikes (referred to as dSpikes) were dependent on dendritic Na^+^ channels and varied in amplitude (Kalmbach et al., 2017).

**Figure 4.**
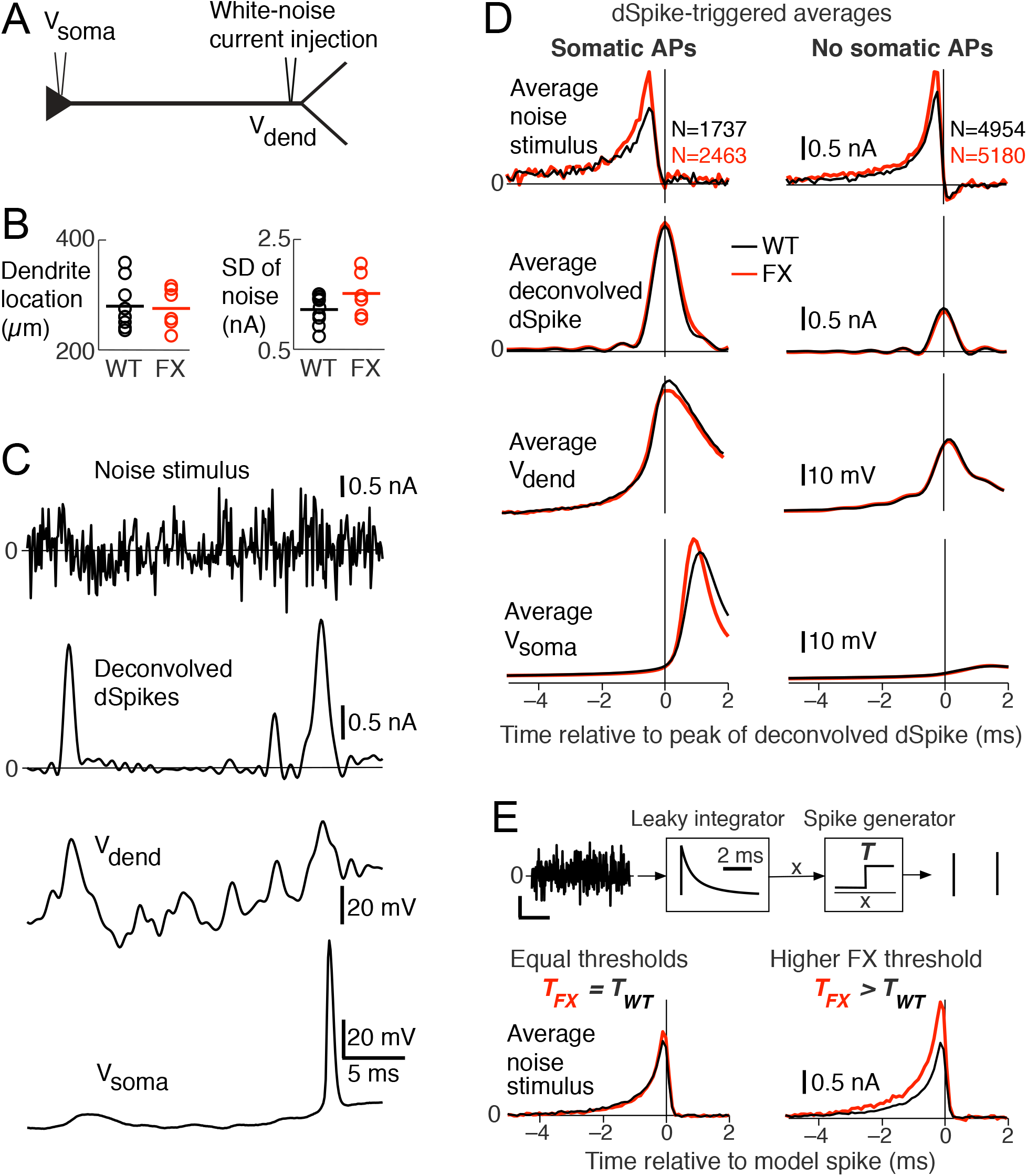
White-noise analysis reveals a higher dendritic spike threshold for FX dendrites. A) One to two minutes of zero-mean Gaussian-distributed white-noise current was injected into the dendrites while simultaneously recording dendritic (V_dend_) and somatic (V_soma_) membrane potentials. B) Location of the dendritic electrode and standard deviation (SD) of the white-noise stimulus for each recording. C) Example V_dend_ and V_soma_ and the deconvolved current dSpikes (note that positive corresponds to an inward current). Only deconvolved current dSpikes with peak amplitudes greater than 0.5 nA were included in the analysis. As illustrated in this example, somatic APs most always followed large deconvolved dSpikes, but large dSpikes also occurred without subsequent APs. D) The average stimulus, V_dend_ and V_soma_ aligned to the peak of the deconvolved dSpike for WT and FX recordings (dSTAs). To reduce variability, the dSTAs were computed from all recording sessions combined. The left column corresponds to dSTAs computed from deconvolved current dSpikes that preceded somatic APs, while the right column shows dSTAs using the remaining deconvolved current dSpikes with peak amplitudes greater than 0.5 nA. E) A leaky integrate-and-fire (IAF) model was used to generate dSpikes using the same stimulus for the WT and FX recordings (see Methods for model parameters). The WT and FX models both used the same leaky integrator, with either the same threshold (*T*_*FX*_ = *T*_*WT*_, left) or different thresholds (*T*_*FX*_ = 1.64 X *T*_*WT*_, right). STAs were then computed from each model ‘s spike output.

A zero-mean, Gaussian-distributed white-noise current was delivered through the dendritic electrode for one to two minutes. Neither the location of the dendritic electrode or the magnitude of the noise injection was significantly different between WT and FX recordings (p = 0.85 and 0.09, respectively, two sample t-test; Fig 4B). As shown by the example recording in Fig. 4C, the noise stimulus produced both hyperpolarized and depolarized dendritic membrane potentials around rest, and both small and large dendritic current spikes were revealed by the deconvolution. As previously reported, the white-noise stimulus produced somatic action potentials that were usually preceded by large dendritic current spikes (Kalmbach *et al*., 2017).

To reveal the dendritic input that produced somatic action potentials, we averaged the dendritic current injection noise aligned on each deconvolved dendritic current spike that preceded each action potential (referred to as the dSpike-triggered-average or dSTA). To reduce variability, we computed the dSTA by combining thousands of somatic APs recorded from all cells. The resulting dSTA stimulus shows that a somatic AP required a larger dendritic stimulus in FX dendrites compared to wild type (p < 1e-8, two-sample t-test; Fig. 4D, left column). We further observed that the average stimulus that preceded deconvolved dSpikes with no subsequent somatic AP was larger for FX versus WT dendrites (p < 1e-6, two-sample t-test; Fig. 4D, right column). Notably, the magnitude of the average deconvolved dSpike, dendritic and somatic potentials were nearly identical between WT and FX dendrites.

Based on the spike-triggered-average theorem, the shape of the dSTA derived using a white-noise stimulus captures the shape of the dendritic integrator. Thus, the timescale of the dendritic integrator was similar, and relatively fast, for both WT and FX dendrites. As reported above, the dSpike threshold was found to be greater for FX dendrites, and we wondered is this could account for the differences in the stimulus STAs. To illustrate the link between a change in dSpike threshold and the magnitude of the stimulus dSTA, we simulated a simple integrate-and-fire (IAF) model using the same white-noise stimuli used in our recordings (see Methods). We set the parameters of the leaky integrator (shown in Fig. 4E) and threshold (*T*_*WT*_ and *T*_*FX*_) to mimic the dSTAs observed in the data. When the stimuli used during the WT and FX recordings were presented to the same IAF models (ie., identical integrators and thresholds), the STAs were very similar (Fig., 4E, left). When the threshold for FX model was increased by 64%, however, the FX STA increased compared to the WT STA in a similar manner to that observed in the data (Fig. 4E, right). Thus, a change in the STA for the FX model stimulus is consistent with an increase in dendritic spike threshold observed in FX dendrites (Figs 1 and 3).

### Neuronal morphology is not different between wild type and FX neurons

Neuron morphology and dendritic branching has a strong influence the electrotonic decay of membrane potential (Rall, 1962). During recordings, neurons were filled neurobiotin for post-hoc morphological reconstructions (Fig. 5A). There was no significant difference in total dendritic length between wild type and FX ET neurons (Fig. 5B; wild type: 6.3 ± 0.85 mm, n=6; FX: 6.2 ± 0.34 mm, n = 7; unpaired t-test: t=0.1395, p =0.8916). Sholl analyses found no significant difference in dendritic length as a function of distance from the soma (Fig. 5C; two-way ANOVA, main effect of genotype: F_(1, 312)_=1.18e^-14^, p>0.99). There was no significant difference in total number of intersections between wild type and FX ET neurons (Fig. 5D; wild type: 188 ± 20 intersections, n=6; FX: 179 ± 6 intersections, n = 7; unpaired t-test: t=0.4626, p =0.6527). Sholl analyses found no significant difference in the number of intersections as a function of distance from the soma (Fig. 5E; two-way ANOVA, main effect of genotype: F_(1, 312)_=0.00, p>0.99). These results suggest that similar to our observations for Layer 2/3 pyramidal neurons in the PFC and CA1 pyramidal neurons in the hippocampus (Routh *et al*., 2017; Ordemann *et al*., 2021), gross dendritic morphology of L5 ET neurons in the PFC are not different in FX mice.

**Figure 5.**
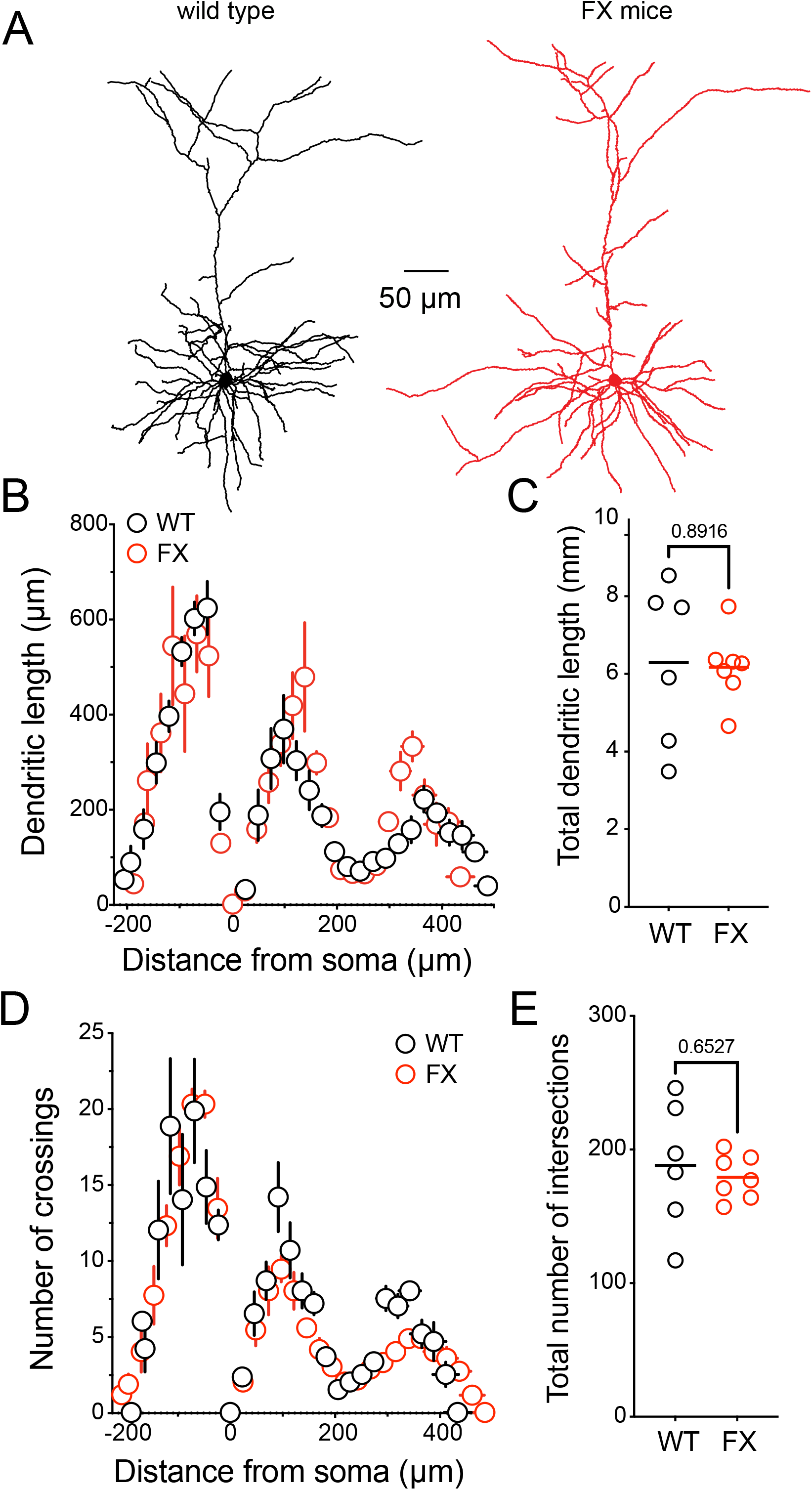
Neuronal morphology is not different between wild type and FX ET neurons. A, Representative Neurolucida reconstructions of ET neurons from wild type (black) and FX (red) mice. B, Sholl analysis of dendritic length is not different between wild type and FX ET neurons. C, There is no significant difference in total dendritic length between wild type and FX ET neurons. D, Sholl analysis of dendritic branching is not different between wild type and FX ET neurons. E, There is no significant difference in total number of intersections between wild type and FX ET neurons.

### Dendritic Na^+^ conductance is lower FX ET neurons

Dendritic Na^+^ and K^+^ channels contribute to the generation and amplitude of dendritic spikes (Golding & Spruston, 1998; Golding *et al*., 1999; Kalmbach *et al*., 2017; Ordemann *et al*., 2021). We previously demonstrated that there were no significant differences in dendritic K^+^ channels between wild type and FX ET neurons (Kalmbach *et al*., 2015). We therefore hypothesized that the more depolarized dendritic spike threshold in FX ET neurons was due to a difference in dendritic Na^+^ channel function. We made outside-out patch clamp recordings of dendritic Na^+^ current from wild type and FX ET neurons using step voltage commands (−80 to +40 mV, Δ10 mV, 50 ms). We found that the peak sodium current in FX dendritic patches was significantly smaller beginning at −30 mV compared to wild type neurons (Fig. 6A-B; two-way repeated measures ANOVA, interaction: F_(24, 180)_ = 11.57, p < 0.0001). The maximum sodium conductance density increased with distance from the soma in both wild type and FX patches. However, the maximum conductance density was lower in FX ET neurons compared wild type (Fig. 6C). Indeed, Na^+^ conductance density was lower in FX ET neurons across the range of membrane potentials tested (Fig. 6D; two-way repeated measures ANOVA, interaction: F_(24, 180)_ = 9.180, p < 0.0001).

**Figure 6.**
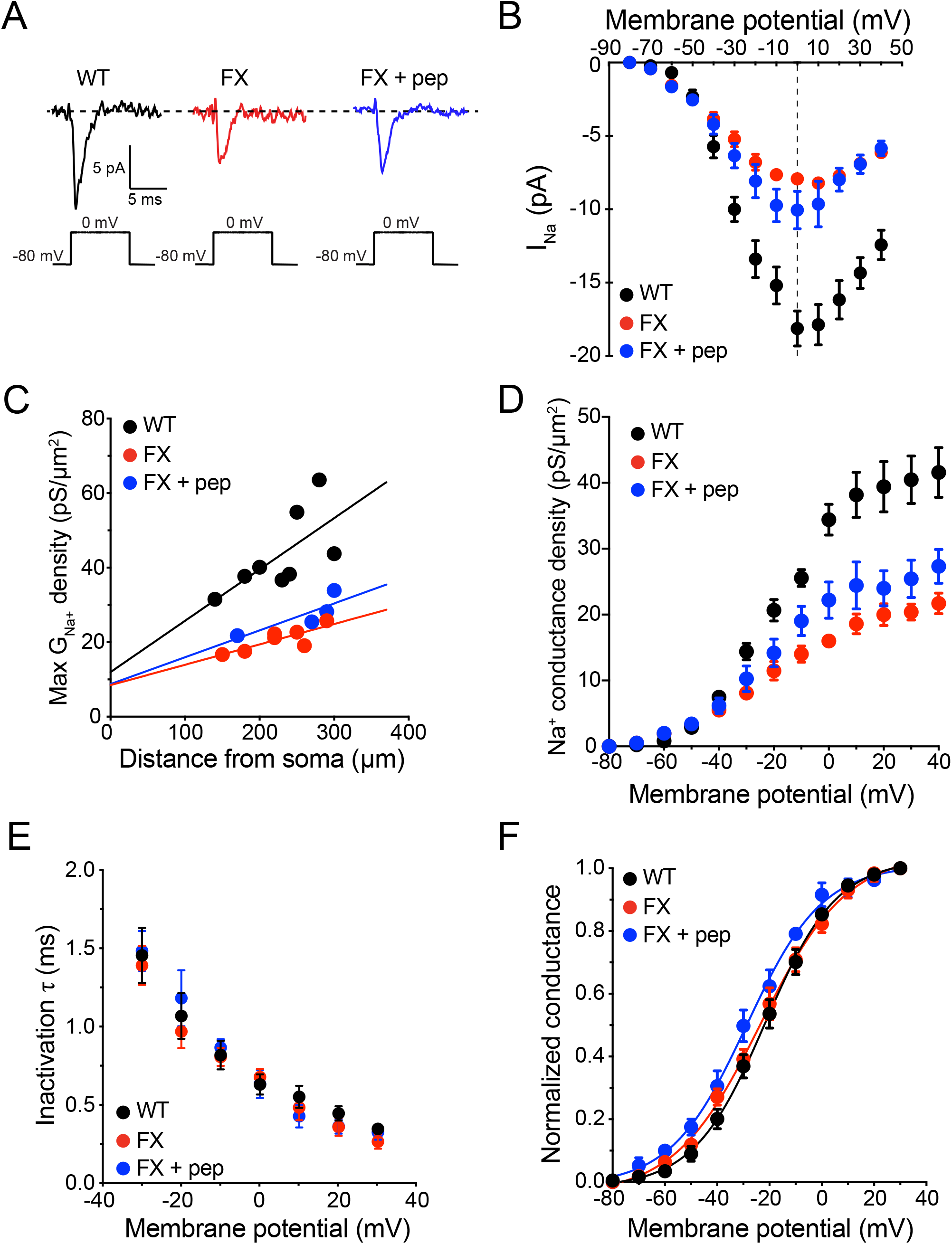
Sodium conductance is lower in FX ET dendrites. A, Representative sodium currents recorded from outside-out patches in response to voltage steps to 0 mV from wild type (black), FX (red), and FX+FMRP_1-298_ peptide (FX + pep, blue) ET dendrites. B, Current-voltage plot showing smaller peak sodium current at more depolarized potentials in FX ET neurons. The presence of FMRP_1-298_ peptide did not rescue peak sodium current. C, The maximum sodium conductance density increases with increasing distance from the soma for both wild type and FX ET neurons but was larger in wild type compared to FX and FX + FMRP_1-298_. D, Summary plot showing lower sodium conductance density as a function of membrane potential in FX and FX + FMRP_1-298_ outside-out dendritic patches. E, Summary plot showing that there was no significant difference in rate of sodium channel inactivation between wild type, FX, and FX + FMRP_1-298_ recordings. F, Summary plot showing that there was no significant difference in sodium channel activation between wild type, FX, and FX + FMRP_1-298_ recordings.

FXS is characterized by the loss of Fragile X Messenger RibonucleoProtein-1 (FMRP; formerly Fragile X Mental Retardation Protein). We previously showed using a truncated FMRP (FMRP_1-298_) that both h-channels and D-type K^+^ channels are regulated by FMRP by protein-protein interactions (Brandalise *et al*., 2020; Kalmbach & Brager, 2020). Intracellular perfusion of FMRP_1-298_ did not however, rescue Na^+^ current or conductance density in outside-out patches from FX ET dendrites (Fig. 6A-D, blue). Consistent with previous results, the rate of Na^+^ channel inactivation increased with membrane depolarization for both wild type and FX dendrites however, there was no significant difference in the rate of Na^+^ channel inactivation between wild type and FX patches (Fig. 6E; two-way repeated measures ANOVA, F_(2, 16)_ = 0.4732, p = 0.6314). The voltage dependence of Na^+^ channel activation was not different between wild type and FX ET dendrites and was not affected by the presence of FMRP_1-298_ (Fig. 6F). There was no significant difference in voltage of half activation (Table 1; one-way ANOVA, F_(2, 16)_ = 1.629, p = 0.2271) or slope factor (Table 1; one-way ANOVA, F_(2, 16)_ = 1.250, p = 0.3130). These results suggest that diminished expression of dendritic Na^+^ channels underlie the depolarized dendritic spike threshold in FX ET neurons.

**Table 1.** Voltage-dependence of sodium channel activation is not different between wild type and FX ET dendrites. Comparison of the voltage of half activation (V1/2) and slope factor of activation curves between wild type, FX, and FX+FMRP_1-298_ outside out patch clamp recordings. P-value is from one-way ANOVA.

|  | wild type | FX | FMRP<sub>1-298</sub> | p-value |
| --- | --- | --- | --- | --- |
| V<sub>1/2</sub> (mV) | -22.4 ± 1.4 | -24.3 ± 1.4 mV | -26.5 ± 1.7 mV | 0.2271 |
| Slope factor (k) | 12.4 ± 0.6 | 13.5 ± 0.6 | 13.7 ± 0.9 | 0.3130 |

## DISCUSSION

Dendritic spikes are critical to information processing in the prefrontal cortex. Distal synaptic inputs onto Layer 5 ET neurons are too weak to overcome the electronic isolation and trigger somatic action potentials. To overcome this, distal synaptic inputs are integrated and transformed into local dendritic Na^+^ spikes. We found that the threshold for dendritic spikes was significantly more depolarized in FX ET neurons compared to wildtype. There was also a smaller proportion of strong dendritic spikes in FX dendrites compared to wild type. Simultaneous recording from the dendrite and soma revealed that dendro-somatic coupling is reduced and that a larger dendritic stimulus is necessary to elicit a dendritic spike in FX dendrites compared to wild type. Lastly, we show that dendritic Na^+^ conductance is lower in FX ET neurons. We suggest that the lower expression of dendritic Na^+^ channels contribute the impaired generation of dendritic spikes in L5 ET neurons of the FX PFC.

### FMRP regulation of Na^+^ channels

Our data suggest that a decrease in Na^+^ conductance (Fig. 6) contributes to the depolarized dendritic spike threshold in L5 ET neurons in the prefrontal cortex of the *Fmr1* KO mouse. FMRP modulates ion channel expression and function through translational mechanisms and through protein-protein interactions (Brager & Johnston, 2014). Several voltage-gated ion channels are regulated by FMRP via protein-protein interactions including BK and SK Ca^2+^-activated K^+^ channels, Na^+^ activated K^+^ channels, D-type K^+^ channels, and HCN-channels (Brown *et al*., 2010; Deng *et al*., 2013; 2019; Brandalise *et al*., 2020; Kalmbach & Brager, 2020). We found no effect of the truncated FMRP on Na^+^ conductance (Fig. 6) suggesting that FMRP does not regulate dendritic Na^+^ channels in L5 ET neurons via protein-protein interactions.

FMRP can both repress and promote translation of target mRNAs (Darnell *et al*., 2011; Banerjee *et al*., 2018). Na^+^ channels in the brain are composed of a pore-forming α subunit (Na_V_1.1, Na_V_1.2, Na_V_1.3, or Na_V_1.6) and auxiliary β subunits (Na_V_β1–4) (Catterall, 2012). Two Na^+^ channel α subunits, Na_V_1.2 (*SCN2A*) and Na_V_1.6 (*SCN8A*), are known mRNA targets of FMRP (Darnell *et al*., 2011). In neurons of the medial prefrontal cortex, reduction of Na_V_1.2 alters dendritic excitability and produces behavioral inflexibility and social impairments consistent with autistic phenotypes (Spratt *et al*., 2019). If FMRP promotes translation of SCN2A or SCN8A, then its loss would decrease the expression of Na_V_1.2 and Na_V_1.6 in L5 neurons respectively. The biophysical properties and surface expression of Na^+^ channel α subunits is regulated in part by β subunits (Patino & Isom, 2010). Accordingly, a change in the expression or association of the Na^+^ channel pore-forming subunit with β subunits could alter the kinetics and/or surface expression of Na^+^ channels. One potential mechanism underlying the lower conductance density in FX neurons would be a decrease in the activation to inactivation balance. We found that the rate of Na^+^ channel inactivation was not different between wild type and FX neurons. While it is possible that the activation rate for FX Na^+^ channels is slower compared to wild type neurons, we could not directly test this hypothesis due to the very rapid activation rate of Na^+^ channels.

### Physiological consequences

We provide the first direct comparison of dendritic spikes between wild type and FX KO neurons in the PFC. We found that the threshold was more depolarized, and the maximum dV/dt was slower for dendritic spikes in FX ET dendrites compared with wild type. We previously showed that dendritic spikes in CA1 pyramidal neurons of the hippocampus also have a depolarized threshold and reduced maximum dV/dt (Ordemann *et al*., 2021). In CA1 dendrites however, there is no difference in dendritic Na^+^ conductance. The change in dendritic spike generation in CA1 neurons is due to a shift in the activation of dendritic A-type K^+^ channels (Routh *et al*., 2013; Ordemann *et al*., 2021). This suggest that although both L5 ET neurons of the PFC and CA1 pyramidal neurons of the hippocampus display the same dendritic spike phenotype, the biophysical mechanisms that underlie this change are different.

The distal dendrites of L5 ET neurons in the PFC receive inputs from the contralateral prefrontal cortex and both the medial dorsal and ventromedial thalamus (Dembrow *et al*., 2015; Collins *et al*., 2018). The dendritic geometry and biophysical properties of the L5 ET dendrites restrict the ability of these distal synaptic inputs to change the somatic membrane potential. Instead, synaptic inputs can be summated to produce local dendritic spikes which are able to reliably propagate to the soma and contribute to action potential firing. In wild type mouse and rat ET neurons, the high density of h-channels in the distal dendrites of L5 ET neurons reduces the window for temporal integration (Kalmbach *et al*., 2013; 2015; Brandalise *et al*., 2020). Accordingly, L5 ET neurons function as coincidence detectors (Dembrow *et al*., 2015). We previously showed that there is a loss of dendritic h-channels and somatic D-type K^+^ channels in FX ET neurons (Kalmbach *et al*., 2015; Brandalise *et al*., 2020; Kalmbach & Brager, 2020). The effect of these changes would increase the window for temporal integration, changing the neuron from a coincidence detector to an integrator, and modulating the threshold for action potential generation. In this study we show that the ability of ET neurons in FX mice to reliably trigger dendritic spikes is impaired due to a loss of dendritic Na^+^ channels. This would effectively reduce the ability of dendritic inputs to trigger somatic action potential output. We suggest that the combined effect of Na^+^, K^+^ and h-channelopathies in Fragile X syndrome impair the processing of information by PFC circuits.

## ADDITIONAL INFORMATION

### Data Availability Statement

The data that support the findings of this study are available upon request from the corresponding author.

### Author Contributions

F.B., B.E.K. and D.H.B. conceptualized and designed the research. F.B. and B.E.K. performed all the experiments. F.B., E.P.C. and B.E.K. analyzed the data. F.B., B.E.K., E.P.C., and D.H.B. interpreted the results of experiments. F.B. and D.H.B. drafted the manuscript. F.B., B.E.K., E.P.C., and D.H.B. revised the final version of the manuscript. All authors have read and approved the final version of this manuscript. All persons designated as authors qualify for authorship, and all those who qualify for authorship are listed.

## Acknowledgments

We thank James Ding for the neuronal reconstructions and Daniel Johnston for critical reading of the manuscript.

